# Automated Multimodal Correlative Registration for Organelle-Specific Molecular Imaging

**DOI:** 10.64898/2026.04.30.721814

**Authors:** Chixiang Lu, Kaiqiang Zhao, Di Cui, Gu Chen, Qian Yang, Hui Yang, Murong Zhao, Kaiyun Song, Mehran Nikan, Zhijie Li, Shanchao Zhao, Jinpeng Cen, Xincheng Qiu, Stephen G. Young, C. Frank Bennett, Punit Seth, Kai Chen, Xiaojuan Qi, Haibo Jiang

**Author notes:** These authors contribute equally (Chixiang Lu, Kaiqiang Zhao).

## Abstract

Mapping subcellular drug distribution is essential for understanding trafficking and off-target effects. NanoSIMS enables chemical imaging of labeled therapeutics, but signal interpretation requires ultrastructural correlation with electron microscopy, a manual and laborious process. We present an automated AI-driven pipeline for correlating chemical and ultrastructural images, enabling multiscale, organelle-precise imaging of molecules in cells and tissues. The method integrates bidirectional optical flow, confidence-guided affine transformation, and automated template matching for cross-scale EM alignment. Morphology-rich ion channels (e.g., ^32^S) estimate transformations that propagate to sparse therapeutic signals (e.g., ^79^Br, ^15^N), overcoming low signal-to-noise challenges. We validate this framework across diverse cell and tissue types, tracking oligonucleotide and antibody therapeutics in vitro and in vivo to reveal cell-type- and organelle-specific distribution patterns. This work establishes a generalizable platform for automated multimodal registration and organelle-resolved subcellular pharmacology.

## Introduction

Understanding the subcellular fate of therapeutic agents—including small molecules, nucleic acid therapeutics, and protein-based biologics—is a fundamental challenge in modern drug development^1–3^. The efficacy and safety of such agents depend on their ability to target specific intracellular compartments (endosomal escape for nucleic acid therapeutics, receptor-mediated trafficking for antibody–drug conjugates, and nuclear or cytoplasmic localization for gene-silencing modalities)^4^. Conversely, off-target accumulation of therapeutic agents in the liver and kidney can drive toxicity and reduce therapeutic windows^5^. Resolving these questions demands spatial tools capable of mapping drug molecules within ultrastructural contexts at nanoscale^6–12^.

NanoSIMS (Nanoscale Secondary Ion Mass Spectrometry) is a powerful technique for spatially profiling elements and isotopes in biological specimens, with lateral resolution down to 50 nm and high sensitivity^13, 14^. By incorporating elemental (*e.g.*, Br) or stable-isotope labels (*e.g.*, ^15^N, ^2^H) into molecules of interest, NanoSIMS can provide semi-quantitative, multiplexed chemical maps of molecular distribution within cells and tissues^7, 9–12, 15–17^. However, NanoSIMS images typically lack sufficient ultrastructural contrast, making it difficult to assign chemical signals to specific organelles or cellular compartments without complementary structural information. Electron microscopy (EM) provides the ultrastructural reference required to interpret NanoSIMS data by resolving membrane-bound organelles, vesicular structures, nuclear architecture, and extracellular matrix with nanometer precision. Correlative electron and ion microscopy (*i.e.*, detecting secondary ions with NanoSIMS) has established the conceptual framework for combining chemical and structural imaging modalities^6, 8, 18–20^. However, the integration of imaging modality information and correlating electron and ion images presents computational challenges, similar to the issues inherent in correlative light and electron microscopy^21–23^. The two modalities differ substantially in contrast mechanisms, spatial scale, field of view, and geometric distortion characteristics, making automated image registration difficult.

Previous approaches to electron and ion image registration have relied largely on manual landmark identification or fiducial marker–based alignment, which are labor-intensive, slow, operator-dependent, and often inconsistent across users and datasets. Semi-automated methods have been proposed, but they require significant user intervention and typically struggle with the low signal-to-noise ratio in NanoSIMS images^20, 24^. Thus far, there is no fully automated, generalizable pipeline that is useful across diverse cell and tissue types and applicable to therapeutic modalities.

Here, we present the CEIM Correlator (Correlator), a fully automated AI-based framework for correlative electron and ion microscopy that enables accurate multimodal registration from tissue scale down to organelle resolution. The key innovation of Correlator is a registration strategy that integrates bidirectional RAFT-based optical flow^25^, confidence-aware affine transformation estimation, and automated cross-scale template matching^26^ to handle the large appearance gap, field-of-view mismatch, and geometric distortion between NanoSIMS and EM images. In addition, by using morphology-rich ion channels to drive registration and transferring the resulting transformation to molecule-specific channels, the method decouples alignment accuracy from the sparse and noisy therapeutic signals that are often the true biological readout. This combination of dense correspondence modeling, reliability-aware transformation fitting, and channel-guided propagation yields a generalizable and user-independent solution for CEIM analysis. We validate the method across multiple cell and tissue types, including bone marrow, liver, spleen, intestine, kidney, and cultured cells, and demonstrate its utility for organelle-level mapping of nucleic acid therapeutics and antibody-based therapeutics *in vitro* and *in vivo*.

## Results

### Overview of the automated CEIM Correlator registration pipeline

We developed a multi-stage computational pipeline for the automated alignment of NanoSIMS and EM images (**Fig. 1**). The core registration strategy is based on bidirectional optical flow estimation using the RAFT deep neural network, which computes dense displacement fields between image pairs. Given an ion image *I*_N_ and an electron micrograph *I*_EM_, RAFT is applied in both the forward direction (*I*_N_ → *I*_EM_) and the reverse direction (*I*_EM_ → *I*_N_), yielding two complementary flow maps: *F*_N→EM_ and ^*F*^EM→N.

**Fig. 1.**
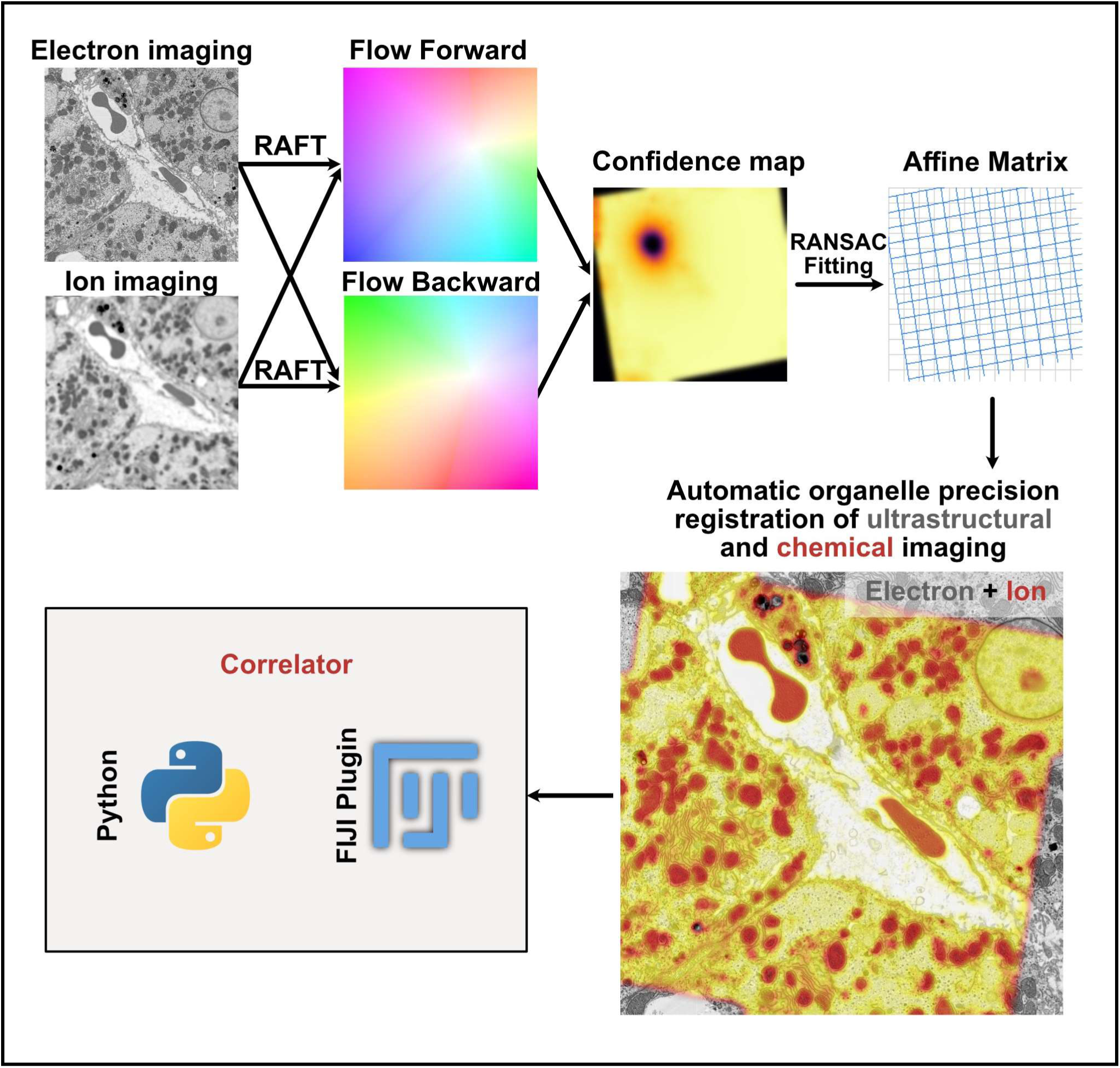
The automated correlative electron and ion microscopy (CEIM) of the Correlator pipeline. Schematic overview of the automated registration pipeline for correlative electron and ion image registration. An ion image and the corresponding EM image are provided as raw inputs. The RAFT (Recurrent All-Pairs Field Transforms) deep neural network is applied in both the forward and backward direction for bidirectional optical flow estimation. A pixel-wise confidence map is computed via forward–backward flow consistency checking and motion-boundary detection. The retained high-confidence point correspondences are used to estimate a global affine transformation matrix. The estimated matrix is applied to warp the full NanoSIMS image stack into the EM coordinate frame using bilinear interpolation, yielding the aligned correlative electron–ion overlay. The code is available in Python, and a Fiji plugin has been developed for use without further coding.

Naive application of all estimated flow vectors for transformation fitting would be unreliable, as optical flow estimation is known to degrade at occluded regions and motion boundaries (where the brightness constancy assumption underlying optical flow breaks down). To identify and exclude such unreliable regions, we employed a two-criterion occlusion detection scheme. First, the forward flow was warped into the target coordinate frame using the backward flow, yielding *F*′_N→EM_pixels where the forward–backward residual magnitude exceeded a relative threshold were flagged as occluded:

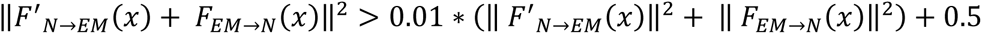

Second, pixels residing near motion boundaries—identified by large spatial gradients in the backward flow—were additionally excluded:

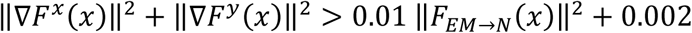

Where ∇*F^x^* is the gradient of forward flow along the *x* axis. The final occlusion map was the logical union of both criteria:

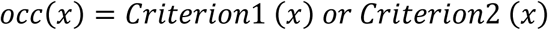

Pixels with *OCC*(*x*) = 0 that are neither occluded nor at motion boundaries were designated as high-confidence correspondences and used exclusively for downstream transformation estimation. In practice, this masking step removed a substantial fraction of flow vectors concentrated at organelle boundaries and image peripheries, where the two modalities exhibit the greatest appearance discrepancy, while retaining the dense, biomass-rich interior regions where morphologically information-rich ion signals and EM contrast are most mutually informative.

The retained high-confidence correspondence points were used to estimate a global affine transformation matrix *A*^∗^ via RANSAC-robust least-squares fitting^27^:

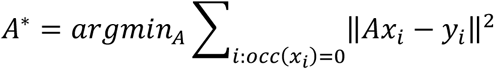

Where *x*_i_ and *y*_i_ are corresponding point coordinates estimated by the network in the electron and ion images, respectively. The affine model accounts for translation, rotation, scaling, and shear—the dominant geometric discrepancies between modalities arising from differences in acquisition geometry, pixel size calibration, and sample preparation. The estimated *A*^∗^is then applied to warp the full ion image stack into the electron micrograph’s coordinate frame via bilinear interpolation (**Fig. 1**). We tested using other ion images (*i.e.*, ^12^C^14^N, ^31^P) that are commonly employed to reveal morphological details in NanoSIMS imaging. The results were similar to using a ^32^S image for registration (**Fig. S1**).

The complete pipeline, from raw image input to aligned multi-channel output, is fully automated, requiring no manual landmark selection, fiducial markers, or user intervention at any stage. The pipeline was implemented in Python, and a Fiji plugin was developed, enabling scientists without programming expertise to use the tool (**Fig. S2**).

### Generalizability of Correlator across cells and tissues

To assess the generalizability of the pipeline across specimen types, we applied Correlator to six different biological specimens: HeLa cells, kidney, bone marrow, intestine, liver, and spleen (**Fig. 2**). These samples encompass a broad spectrum of biological complexity, from single cells to highly organized tissues, and from mineralized tissue to soft organs. For each specimen type, we present the raw EM image, the ^32^S image of selected regions, and the overlay of the “Correlator aligned” ^32^S NanoSIMS on the EM image.

**Fig. 2.**
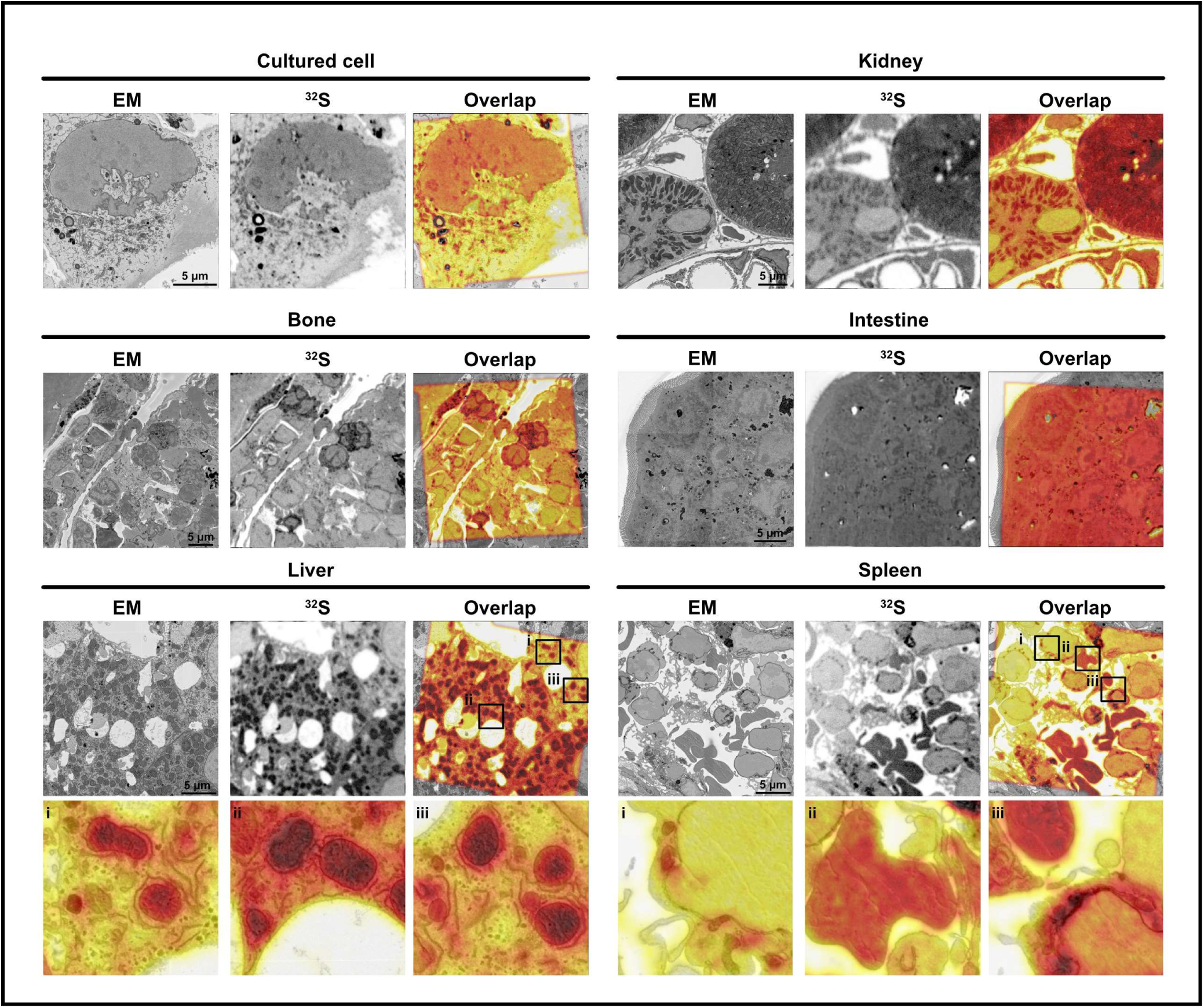
Generalizability of the Correlator across diverse biological specimens. Demonstration of Correlator with six specimens: HeLa cells, kidney, bone marrow, intestine, liver, and spleen. For each specimen, three panels are shown from left to right: (i) raw EM image providing ultrastructural context; (ii) the corresponding ^32^S NanoSIMS image reflecting the distribution of sulfur-containing biomolecules as a proxy for biomass; and (iii) pseudocolor overlay of the registered ^32^S and EM images, demonstrating precision registration of chemical and ultrastructural features.

Across all six specimen types, the pipeline achieved visually accurate alignment, with ^32^S secondary ion signals localizing with the corresponding ultrastructural features in the EM images. The pipeline performed robustly despite variation in tissue morphology, cell heterogeneity, and image quality and resolution across specimen types, demonstrating that the optical flow approach generalizes beyond the tissue context in which it was developed.

### Large-field EM alignment via template matching for cross-scale registration

In many experiments, EM images are captured as large mosaic maps of tissue samples that cover areas larger than the NanoSIMS field of view. The large EM tissue mosaics often help guide the selection of areas for NanoSIMS imaging. In these cases, the first step of the pipeline—optical flow estimation between the full EM and NanoSIMS images—is not directly applicable due to the size mismatch and the need to first localize the NanoSIMS acquisition region precisely within the EM mosaic. We further developed an automatic two-stage alignment strategy to address this issue (**Fig. 3a**).

**Fig. 3.**
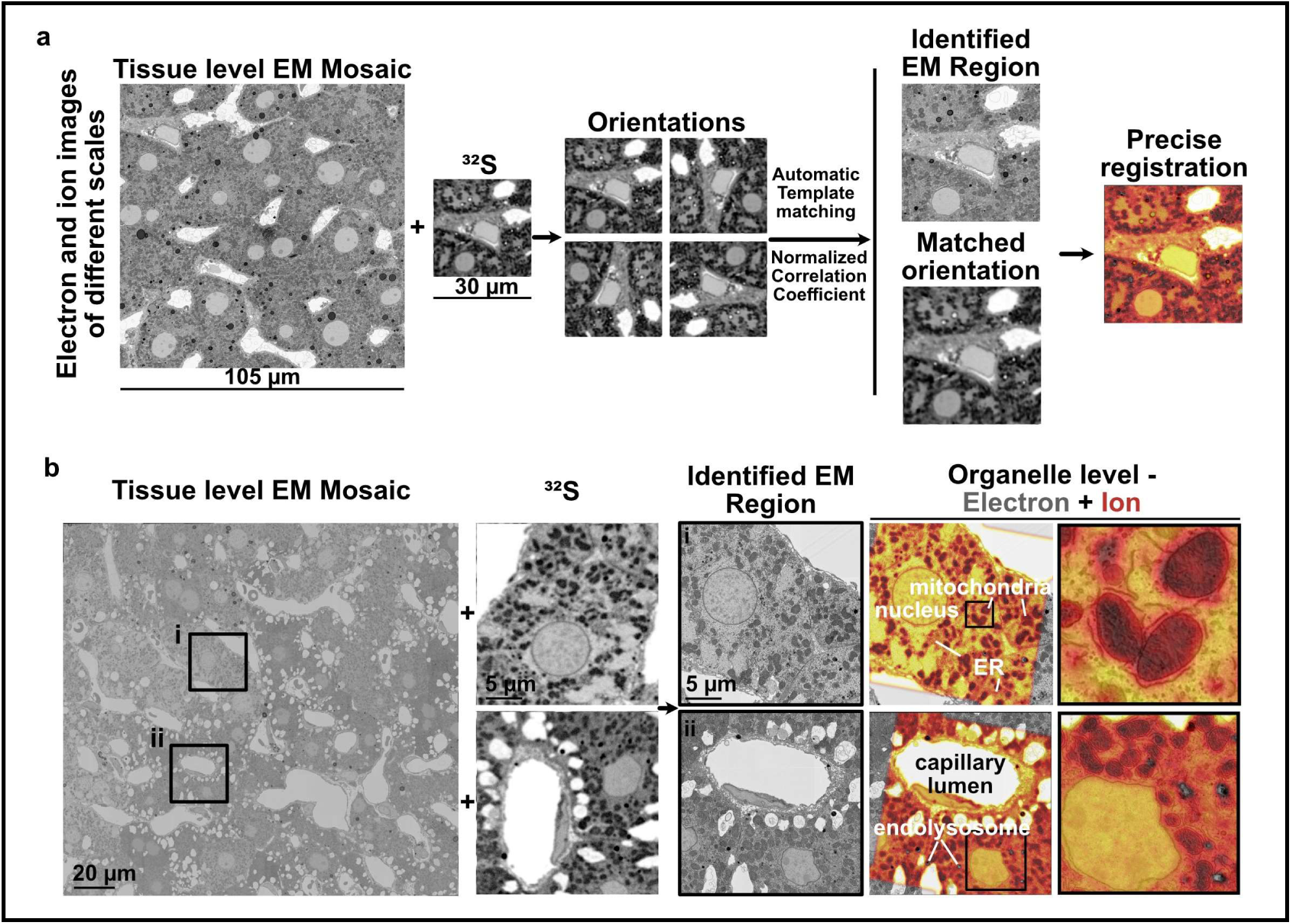
Large-field EM alignment via template matching for cross-scale registration. (**a**) Schematic of the two-stage alignment strategy for large-field EM mosaic datasets. The target ion image is evaluated across all eight canonical orientations (four 90° increment rotations combined with horizontal mirroring) against the large tissue-level EM mosaic using normalized cross-correlation. The coarse EM crop identified via template matching is passed to the RAFT-based bidirectional optical flow pipeline to estimate the affine transformation matrix. (**b**) Application to a large-field liver EM mosaic containing two spatially distinct ion imaging regions (Region 1 and Region 2). For each region, the ^32^S image, identified coarse EM patch, and the overlap of ^32^S ion and EM images are shown.

The secondary ion image template was systematically evaluated across all eight canonical orientations — formed by combining four 90°-increment rotations with horizontal mirroring — against the resolution-matched EM mosaic with normalized cross-correlation (NCC). The orientation and location that jointly yielded the highest NCC score were selected, thereby recovering both the position and the geometric relationship of the ion images within the EM mosaic (**Fig. 3a**).

The coarse EM crop identified by template matching was passed to the RAFT-based optical flow registration pipeline (Sections 2.1 and 4.4.2–4.4.4), which computed the precise affine transformation between the best-orientation secondary ion image and the extracted coarse EM crop. This two-stage design separates the global search problem (handled efficiently by NCC over a discrete set of orientations) from the local sub-pixel refinement problem (handled by RAFT optical flow), yielding accurate alignments regardless of the initial orientation of the image acquisitions.

We applied this pipeline to a large-field liver EM mosaic with ^32^S and ^31^P ion images from two separate regions (**Fig. 3b** and **Fig. S3**). For both cases, the orientation-exhaustive template matching identified the ion image field of view within the mosaic and recovered the correct orientation, as confirmed by the high NCC scores (> 0.8) and visual inspection of the alignments. Subsequent optical flow refinement yielded precise organelle-level registration. The aligned overlays of ^32^S, ^31^P, and EM images revealed the distribution of various organelles, for example the sulfur-enriched endosomes and phosphorus-rich nuclei in hepatocytes, demonstrating the pipeline’s capacity to correlate chemical data and ultrastructural images across different scales.

### Morphology channel-guided correlation for organelle-level visualization of element-labeled nucleic acid therapeutics

An advantage of NanoSIMS chemical imaging is its ability to simultaneously acquire multiple secondary ion signals, enabling co-localization of structural markers and signals for molecules of interest within a single acquisition. However, molecule-specific channels [*e.g.*, ^79^Br for Br-labeled antisense oligonucleotides (ASOs)] often exhibit sparse, punctate signals that are insufficient for robust image registration (**Fig. S4**). To address this issue, we developed a morphology and secondary ion–guided registration strategy in which secondary ions rich in morphological information (*e.g.*, ^32^S) are used exclusively for transformation estimation, and the resulting transformation is then applied to all other ion channels (**Fig. 4a** and **Fig. S4**). This approach is valid because the secondary ion channels are acquired simultaneously and are therefore in perfect registration, such that the geometric transformation correlating NanoSIMS and EM images is channel independent. NanoSIMS images that were acquired with multiple frames were pre-processed with NanoSIMS Stabilizer^28^.

**Fig. 4.**
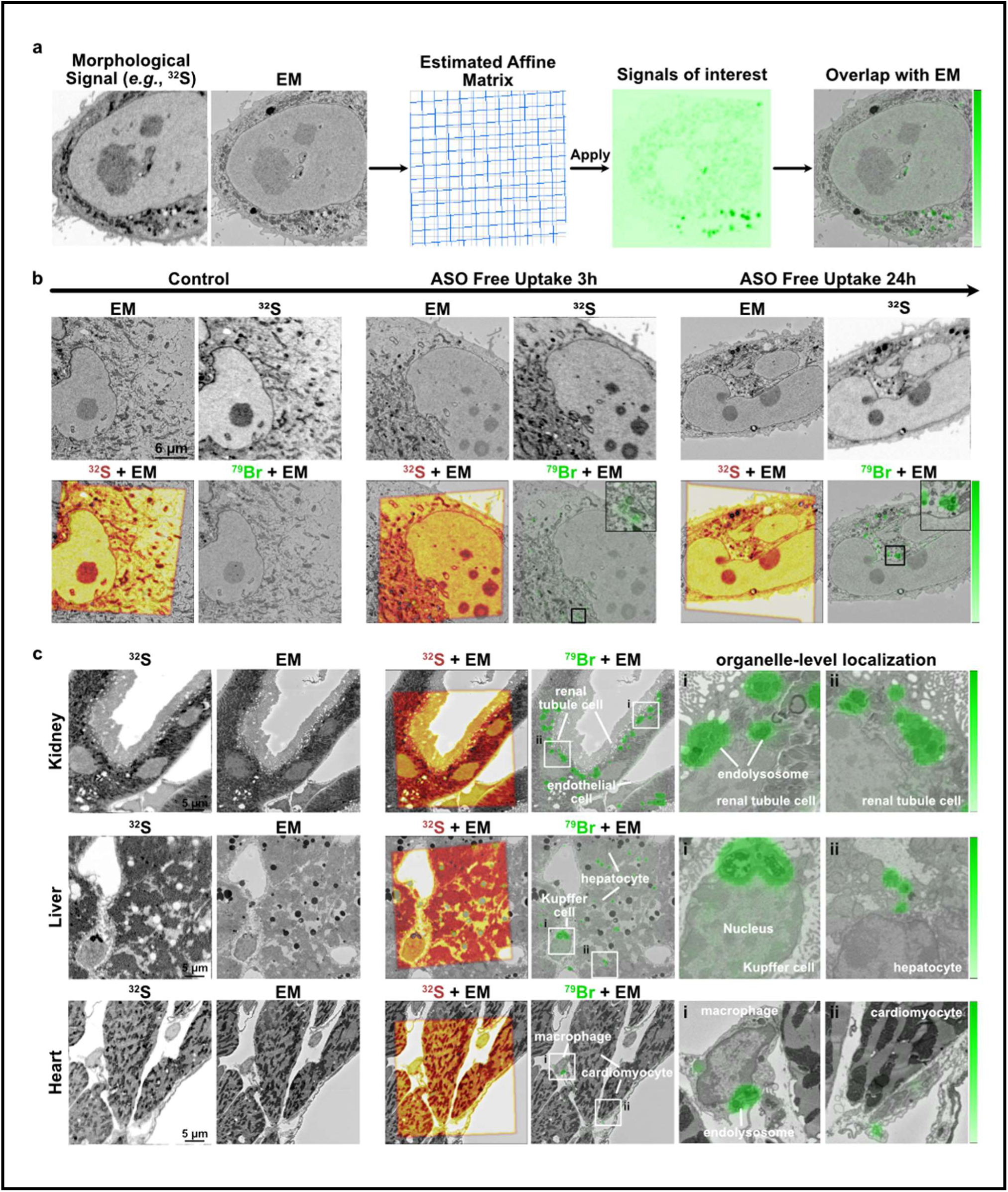
Morphology channel-guided correlation for visualizing the nucleic acid therapeutics. (**a**) Schematic of the morphology channel-guided registration strategy for multi-channel NanoSIMS data. The morphology and contrast-rich channels, for example the ^32^S image, are used to estimate the affine transformation relating ion to EM coordinates. The derived transformation is subsequently propagated to channels that detect isotopes of interest (*e.g*., ^79^Br, ^15^N), thereby decoupling registration accuracy from potentially sparse signals from these isotopes. (**b**) Time-resolved subcellular tracking of Br-labeled antisense oligonucleotide (ASO) uptake in HeLa cells at 0, 3, and 24 h post-treatment. For each time point, four panels are shown: raw EM image; ^32^S image; ^32^S and EM image overlay; and ^79^Br and EM image overlay. (**c**) Organelle-level mapping of ^79^Br-labeled ASOs in kidney, liver, and heart after treatment with 32 mg/kg ASO. Six panels are shown: ^32^S image; EM image; overlay of aligned ^32^S and EM image; overlay of aligned ^79^Br and EM image; two cropped Zoom-in regions showing the distribution of ASOs in organelles of different types of cells.

We demonstrate the pipeline’s capability to track molecules-of-interest, for example oligonucleotide therapeutics, at the subcellular level. We first analyzed HeLa cells that were either untreated or had undergone treatment with Br-labeled ASOs (5 µM) for 3 and 24 h. Following ASO treatment, ^79^Br signals in cells were detected within 3 h and increased further by 24 h, progressively concentrated in intracellular compartments, largely endolysosomes, with a small portion escaping from endolysosomes and accumulating in the nucleus. The aligned chemical and ultrastructural overlays enabled spatial visualization of ASO accumulation over time. Together, these time-resolved data demonstrate the pipeline’s capability to follow dynamic drug partitioning at organelle resolution (**Fig. 4b**).

Next, we applied the multi-channel pipeline to track ASOs in tissues, including kidney, liver, and heart sections from mice that had received Br-labeled nucleic acid therapeutics (**Fig. 4c** and **Fig. S5**). Following ^32^S-guided alignment, the ^79^Br signal was mapped onto the EM ultrastructure, revealing the location of the therapeutic agent in tissues at the single-cell and single-organelle resolution. The alignment of ^79^Br and EM images revealed cellular heterogeneity in drug distribution, with distinct cell populations and subcellular compartments having different levels and patterns of drug accumulation. For example, ASO accumulation in the heart was enriched in macrophages, particularly in endolysosomal compartments, with lower levels of accumulation in cardiomyocytes. In the liver, Kupffer cells exhibited substantial accumulation of ASOs in endolysosomes, along with greater accumulation in the nucleus. These findings highlight the importance of multiscale analysis, from tissues to organelles, for understanding cell type–specific uptake and subcellular fate of therapeutic agents.

### Correlator for organelle-scale visualization of stable isotope–labeled nanobodies in vitro and in vivo

To demonstrate the broad application of Correlator for subcellular drug mapping, we applied it to ^15^N-labeled antibody therapeutics to define the subcellular fate of B12-PE38, a nanobody-based immunotoxin for aggressive variant prostate cancer (AVPC^29^). B12-PE38 consists of a high-affinity nanobody (B12) specific for CEACAM5, a cell-surface antigen (upregulated in AVPC^30, 31^), fused to a truncated *Pseudomonas* exotoxin (PE38). B12-PE38 undergoes receptor-mediated internalization, delivering PE38 payload intracellularly to inhibit protein synthesis and trigger apoptosis^32, 33^ in CEACAM5-positive tumor cells^29^. Resolving the organelle-scale distribution of such therapeutic agents is essential for understanding their intracellular trafficking, informing pharmacokinetic-pharmacodynamic (PK-PD) relationships, and guiding strategies to minimize off-target uptake and systemic toxicity. Using Correlator, we performed organelle-scale mapping of ^15^N-labeled B12-PE38 to resolve its intracellular fate in cultured PC3 prostate cancer cells *in vitro* and after systemic administration *in vivo*.

We first applied Correlator to map the subcellular distribution of [^15^N]B12-PE38 in cultured PC3 prostate cancer cells. PC3 cells were treated with [^15^N]B12-PE38 and processed for CEIM imaging. Following automated alignment, the ^15^N signal was mapped onto corresponding EM images (**Fig. 5a**), revealing ^15^N enrichment in endolysosomes and endoplasmic reticulum (ER), consistent with the intracellular trafficking pathway of PE38-based immunotoxins^34^. To delineate the subcellular fate of B12-PE38 in a physiologically relevant context, we extended our analyses to mice with PC3 xenografts. Previous studies established that B12-PE38 accumulates in the tumor following systemic administration and is cleared predominantly by liver and kidney^29^. We focused our analysis on the xenografts and those tissues to visualize on-target delivery and off-organ clearance at the organelle level. Mice bearing PC3-derived xenografts received a single tail vein injection of [^15^N]B12-PE38. Tumor, liver, and kidney were collected, processed, imaged by CEIM, and analyzed with Correlator. In tumor tissue, the ^15^N signal was detected in the ER (**Fig. 5b** and **Fig. S6a**), corroborating the requirement of retrograde transport for PE38-mediated protein synthesis inhibition^34^. We also observed an elevated ^15^N signal in the nucleus of tumor cells (**Fig. 5b** and **Fig. S6a**). This observation suggests that some of B12-PE38 undergoes nuclear import, but further studies are warranted. ^15^N accumulation was observed in erythrocytes and in endolysosomes and ER of liver and kidney (**Figure 5c-d** and **Fig. S6b-c**), consistent with the observation that B12-PE38 is cleared by the liver and kidney^29^. Interestingly, we also detected ^15^N signal in the mitochondria of hepatocytes and renal tubule cells (**Fig. 5c-d** and **Fig. S6b-c**). Given that PE38 has been reported to induce apoptosis via mechanisms involving mitochondrial damage^35, 36^, we speculate that mitochondrial association of B12-PE38 or its fragments may contribute to its cytotoxic activity, although the functional significance of the mitochondrial localization remains unclear. Together, these tissue-level data provide a quantitative, organelle-level mapping of B12-PE38 distribution across both the tumor itself and the clearance organs, providing useful information for assessing on-target efficacy and off-tissue toxicity.

**Fig. 5.**
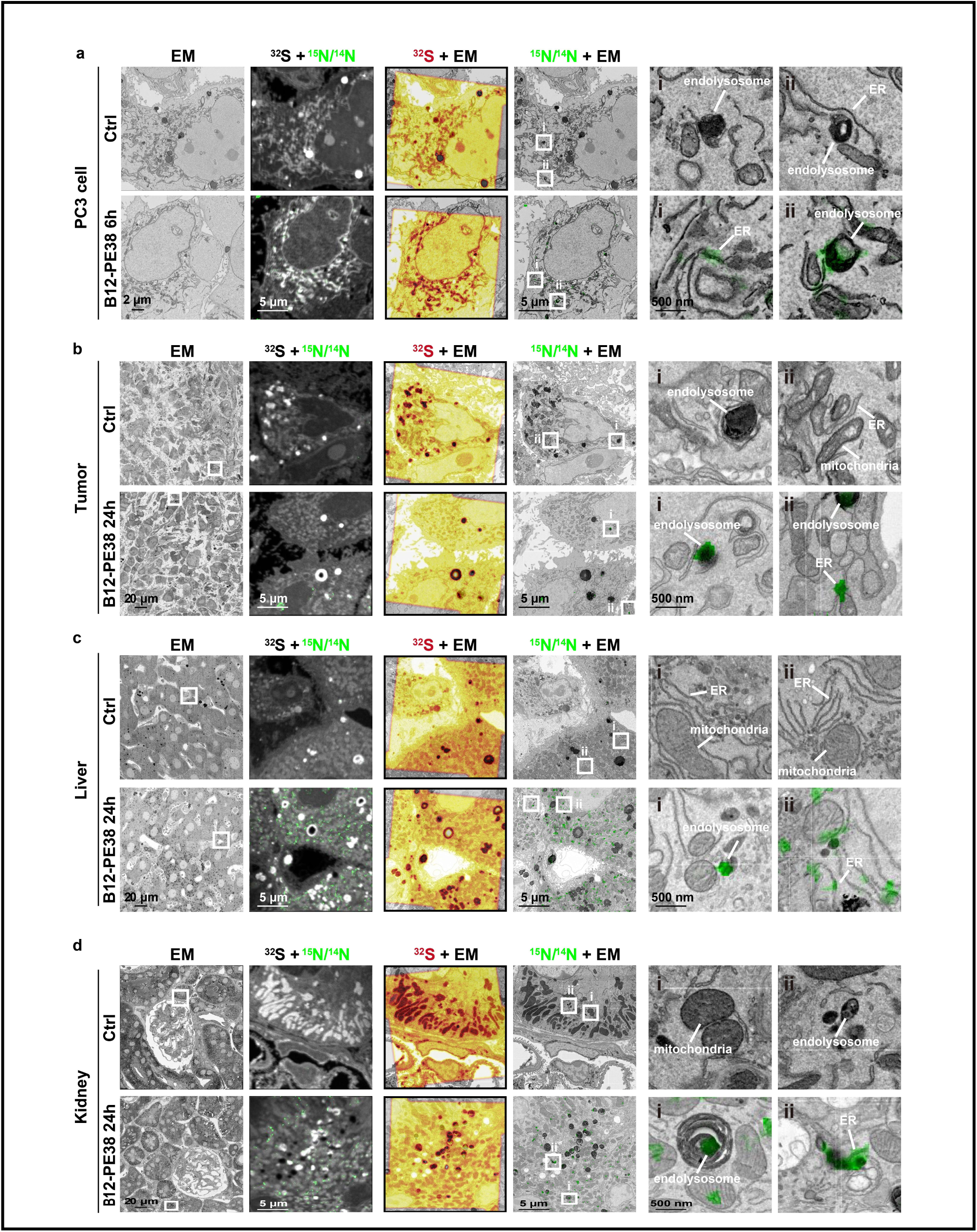
Organelle-level mapping of B12-PE38 distribution *in vitro* and *in vivo*. Subcellular visualization of [^15^N]B12-PE38 in PC3 cells 6 h post-treatment (**a**) and in tumor (**b**), liver (**c**), and kidney (**d**) 24 h after administration. From left to right: EM image; ^15^N/^14^N and ^32^S overlay image; ^32^S and EM overlay; ^15^N/^14^N and EM overlay; two representative Zoom-in regions from ¹⁵N/^14^N and EM overlays showing organelle-level distributions of B12-PE38.

## Discussion

The automated CEIM Correlator pipeline addresses a key bottleneck in multiscale and multimodal correlative image analysis, achieving robust, fully automated alignment across diverse biological specimens without manual and potentially biased landmark selection. By integrating chemical information from NanoSIMS images with ultrastructural features of electron micrographs, this workflow provides organelle-level precision localization across scales and biological contexts. We demonstrate that this capability is important for studying subcellular drug partitioning.

The key innovative feature of Correlator is the use of bidirectional optical flow to construct a pixel-wise, occlusion-based confidence map that identifies correspondence points despite differences in resolution and contrast between chemical and ultrastructural images. This locally adaptive strategy is more robust in the setting of heterogeneous contrast and noise characteristics of NanoSIMS data than global intensity–based registration methods. Equally important is the structured channel-guided registration strategy, in which the morphologically informative ion channels (*e.g.*, ^32^S) drive transformation estimation while sharing the alignment with molecular-specific ion channels. This decoupling ensures that reliable registration is independent of the sparse, low-SNR signals characteristic of ion channels for molecules of interest. For tissue-level large-field EM mosaics, the orientation template matching stage extends the pipeline’s applicability to datasets in which the NanoSIMS field of view is a small fraction of the total EM area and the relative acquisition orientation is unknown—scenarios that are increasingly common with automated EM platforms.

Beyond its technical performance, the pipeline demonstrated important applications of organelle-scale mapping of therapeutic partitioning. In our studies, the workflow resolved drug localization not only across different tissues, but also across distinct cell populations and subcellular compartments. This feature is particularly valuable for identifying cell-to-cell heterogeneity in therapeutic uptake and intracellular fate, which is obscured by bulk tissue measurements or by fluorescence-based imaging approaches with limited resolution. Applications to ASOs and B12-PE38 nanobody distribution in cells and tissues illustrate the translational value of the approach, providing organelle-scale drug localization with specificity not attainable with fluorescence microscopy.

Several limitations remain. The RAFT network was trained on natural image benchmarks; domain-specific fine-tuning could further improve performance with low-SNR or morphologically unusual specimens. Also, the pipeline assumes co-registration of NanoSIMS and EM on the same section; the ^133^Cs^+^ sputtering that is required for NanoSIMS imaging removes the surface layer, potentially resulting in inaccurate registration between EM and NanoSIMS images. Despite these limitations, the current studies establish a generalizable and scalable framework for automated correlative chemical and ultrastructural imaging. We foresee that similar strategies could be used for correlative analysis of other imaging modalities.

## Code Availability

Source code of Python, FIJI plugin, and documents are available in https://github.com/Luchixiang/Correlator

## Acknowledgments

This work is supported by the Innovation and Technology Fund (MHP/129/22), Hong Kong Research Grant Council General Research Fund (17102722, 17300523, 17302324), National Natural Science Foundation of China (32271445) to H.J; Australian Research Council (ARC) Early Career Industry Fellowship (IE230100042) and the National Health and Medical Research Council (NHMRC), Australia, Ideas Grants (2038782) to K.C.; and a grant from the Leducq Foundation (23CVD02) to S.G.Y. The work was conducted in the JC STEM Lab of Molecular Imaging, funded by The Hong Kong Jockey Club Charities Trust.

## Author contributions

HJ, CL, and KZ designed the experiments and wrote the paper. CL, KZ, DC, GC, QY, HY, KS, XQ, MZ, JC, ZL, SZ, PS, CB, MN, SGY, and KC performed experiments, collected, analyzed, and assembled the results. HJ, SGY, and KC secured funding. All have commented on and edited the manuscript.

## Materials and Methods

### Purification of ¹⁵N-labeled B12-PE38 conjugate

The expression and purification of ^15^N-labeled B12-PE38 conjugate were performed as previous described^29^, with the following modifications for isotopic labeling. *Escherichia coli* BL21 (DE3) cells harboring the pET-22B-B12-PE38 plasmid were grown in 1 L of LB medium at 37 °C until the OD₆₀₀ reached 1.0. The cells were then harvested by centrifugation (4,000 × g, 20 min), washed twice with sterile M9 minimal medium lacking a nitrogen source. One-third of the bacterial pellet was transferred into 1 L of M9 minimal medium supplemented with ¹⁵N-labeled ammonium chloride (¹⁵NH₄Cl, 1 g/L) as the sole nitrogen source, along with 4 g/L D-glucose, 2 mM MgSO₄, 0.1 mM CaCl₂, and appropriate antibiotics. The culture was incubated at 37 °C with shaking until OD₆₀₀ reached 0.8, and protein expression was induced by the addition of isopropyl β-D-1-thiogalactopyranoside (IPTG) to a final concentration of 1 mM and the culture was grown overnight (over 18 h) at 30 °C. Cells were harvested by centrifugation (8,000 × g, 20 min) and the subsequent purification steps were carried out as described previously^29^.

### Cell lines and cell culture

HeLa and PC3 cell lines were obtained from American Type Culture Collection (ATCC), Manassas, VA, USA and maintained in α-minimum essential medium (HeLa) or Ham’s F-12 K medium (PC3) supplemented with 10% fetal bovine serum (FBS), 100 U/mL penicillin, and 100 μg/mL streptomycin. For ASO treatment experiments, ASOs were synthesized by Ionis Pharmaceuticals and labeled as we previously described^9^. HeLa cells were incubated with PBS or 5 μM bromine-labeled ASOs for 3 or 24 h. For ¹⁵N-labeled B12-PE38 treatment experiments, PC3 cells were incubated with PBS or 20 μg/mL ¹⁵N-labeled B12-PE38 for 6 h. Both cells were maintained at 37 °C in a humidified atmosphere containing 5% CO₂.

### Preparation of cultured cells for correlative electron and ion microscopy

After treatment, cells were fixed for 2 h in prewarmed 2.5% glutaraldehyde prepared in 0.1 M sodium cacodylate buffer (pH 7.4). Cells were then washed five times for 3 min each in ice-cold 0.1 M sodium cacodylate buffer. Post-fixation was carried out on ice for 60 min using 2% osmium tetroxide and 1.5% potassium ferricyanide in 0.1 M cacodylate buffer while cells remained on coverslips. After washing, coverslips were transferred to a fresh 24-well plate and incubated in 1% thiocarbohydrazide for 30 min at room temperature. Samples were then washed again and treated with 2% osmium tetroxide in water for 30 min at room temperature. Following additional washes, cells were incubated overnight at 4°C in 2% uranyl acetate or niobium acetate. After staining, cells were dehydrated through sequential incubations in 30%, 50%, 70%, 90%, and 100% ethanol, followed by 100% acetone, for 3 min at each step. Resin infiltration was performed using 33%, 50%, 75%, and 100% EMbed 812 in acetone for 2 h per step, followed by overnight incubation in 100% resin. Coverslips were then inverted onto resin-filled BEEM capsules with the cell layer facing the resin and polymerized at 60°C for 48 h. To detach the glass, polymerized blocks were inverted, briefly immersed in liquid nitrogen for 15 s, and then returned to room temperature for 10 s. Once the coverslip surface turned white, it was removed with tweezers, leaving the embedded cell monolayer on the resin surface. Blocks were trimmed and sectioned at 500 nm thickness using a Leica ultramicrotome and a Diatome diamond knife. Sections were collected on silicon wafers for subsequent imaging.

### Animals and Ethical Statement

For ASO injection experiments, 4-month-old male C57BL6/J mice were obtained from Animal Resources Centre (Canning Vale, WA, Australia). Animal care and experiments were done in accordance with Animal Welfare Act 2002 (WA) and the Australian Code for the Care and Use of Animals for Scientific Purposes (eighth edition, 2013). Procedures involving animals were approved by The University of Western Australia Animal Ethics Committee (Approval No. 2021/ET00075). For ^15^N-labeled B12-PE38 injection experiments, Nude mice (6–8 weeks old) purchased from GemPharmatech (Guangzhou, China). All animal procedures were conducted under a protocol approved by the Institutional Animal Care and Use Committee (IACUC) of the Shenzhen People’s Hospital and in accordance with relevant institutional and national guidelines (Approval No. AUP-230224-LZJ-543-01).

### Animal experiments

For ASO tracing, 4-month-old male C57BL6/J mice were intraperitoneally injected with 32 mg/kg bromine-labeled ASOs. After 24 h, the animals were euthanized, and liver, heart, and kidney tissues were harvested and immediately processed for correlative electron and ion microscopy sample preparation. For B12-PE38 tracing experiments, Nude mice (6–8 weeks old) were subcutaneously inoculated in the right flank with PC3 human prostate cancer cells (5 × 10⁶ cells in 100 μL of PBS). Tumor size was measured with vernier calipers and calculated using the following formula: (length × width × width)/2. When tumor volumes reached approximately 150 mm³ (n = 3 per group), mice were administered a single tail vein injection of either sterile PBS (control) or 0.6 mg/kg ¹⁵N-labeled B12-PE38 conjugate. After 24 h, the animals were euthanized, and tumor, liver, and kidney tissues were harvested and immediately processed for correlative electron and ion microscopy sample preparation.

### Animal euthanasia and tissue harvest

Following treatment, mice were euthanized by cervical dislocation, and heart perfusion was performed through the left ventricle using ice-cold heparinized saline (10 U/mL; Sigma), followed by freshly prepared fixative containing 2% paraformaldehyde, 2.5% glutaraldehyde, and 2.1% sucrose in 0.1 M sodium cacodylate buffer (pH 7.4). Tissues of interest were then dissected, trimmed into smaller pieces, and immersion-fixed in the same solution at 4°C for 3 days with gentle rotation. Fixed samples were subsequently rinsed five times in ice-cold 0.1 M sodium cacodylate buffer for 5 min per wash.

### Preparation of tissue samples for correlative electron and ion microscopy

Following fixation and buffer washes, all tissue samples were first incubated on ice for 60 min in 2% osmium tetroxide and 1.5% potassium ferricyanide prepared in 0.1 M cacodylate buffer. Samples were then washed five times in cold 0.1 M cacodylate buffer (5 min each) and treated with 1% thiocarbohydrazide for 30 min at room temperature. After extensive washing in water, specimens were exposed to 2% osmium tetroxide for 1 h at room temperature, washed again, and then incubated overnight at 4°C in 2% uranyl acetate or niobium acetate. The following day, samples were washed and dehydrated through a graded ethanol series (30%, 50%, 70%, 90%, 100%, 100%), followed by 100% acetone, with each step lasting 5 min. Resin infiltration was then performed stepwise using EMbed 812 resin diluted in acetone at 33%, 50%, 75%, and 100% concentrations for 2 h each, followed by overnight incubation in pure resin. Samples were polymerized in fresh 100% EMbed 812 resin at 60°C under vacuum for 48 h. Hardened blocks were trimmed and sectioned at 500 nm thickness using a Leica ultramicrotome equipped with a Diatome diamond knife. Sections were collected on silicon wafers for subsequent imaging.

### Scanning electron microscopy

For tissues and cultured cells in Figures 1-3, 4c, 5, S1, S3, and S6, backscattered electron (BSE) images were acquired using a GeminiSEM 360 scanning electron microscope (ZEISS). Resin sections from both tissues and cultured cells in Figure 4a-b and Figure S4-5 were imaged using an FEI Verios scanning electron microscope (Thermo Fisher Scientific). Backscattered electron detection was used to generate ultrastructural images.

### NanoSIMS imaging

Sections were coated with a 5 nm layer of gold and analyzed using either a NanoSIMS 50 or NanoSIMS 50L instrument (CAMECA, France). The regions of interest were scanned with a focused ^133^Cs^+^ primary ion beam operated at 16 keV, and secondary ions (*e.g.*, ^32^S, ^31^P, ^79^Br, ^12^C^14^N) were collected using multicollection detectors to generate ion images. Before image acquisition, selected regions were subjected to a presputtering step with ^133^Cs^+^ implantation to remove the surface gold coat and stabilize secondary ion emission. Presputtering was performed using a primary beam current of approximately 1 nA with a D1 = 1 aperture until a dose of about 1 × 10^17^ ions/cm^2^ was reached for ASO imaging, and approximately 5 × 10^16^ ions/cm^2^ dose for B12-PE38 imaging. For imaging, scans were acquired using a D1 = 3 aperture with a primary current of ∼2 pA.

### CEIM Correlator Pipeline

#### Image pre-processing

First, NanoSIMS images with multiple frames were aligned with NanoSIMS Stabilizer^28^. Then, prior to registration, NanoSIMS and EM images were preprocessed to normalize intensity distributions and reduce noise. NanoSIMS images were smoothed with a Gaussian filter (σ = 3 pixel) and intensity-normalized to the range [0, 1]. EM images were normalized to [0, 1]. EM images were down-sampled to a NanoSIMS image pixel size using bilinear interpolation.

#### Bidirectional Optical Flow Estimation

The optical flow was calculated using the RAFT algorithm. Similarities between the two feature maps were measured:

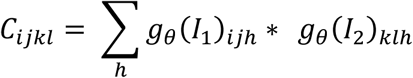

Where *g*_0_is the feature extractor and I represents the image. The similarity was used to guide the optical flow map estimation in GRU.

RAFT was applied in both the forward direction (NanoSIMS → EM) and the reverse direction (EM → NanoSIMS), yielding flow fields *F*_N→EM_, and *F*_EM→N_.

Of note, our experiments revealed that the RAFT weight provided by the author of RAT, pre-trained on the FlyingThings dataset, is capable of estimating the optical flow in NanoSIMS images. Consequently, there is no necessity to add additional training or fine-tuning of the RAFT algorithm.

#### Large-Field Template Matching

Eight orientation variants of the preprocessed NanoSIMS image were generated by applying all combinations of four 90°-increment rotations and horizontal mirroring. For each invariant *I*^K^ normalized cross-correlation template matching was performed against the pixel size-normalized EM mosaic.

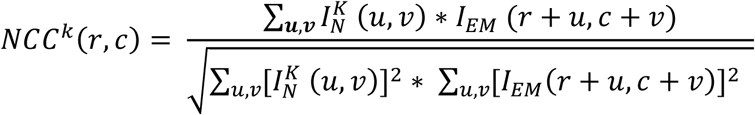

The best-matching orientation and location were jointly selected as:

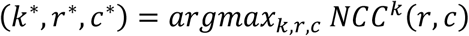

The orientation label k, encoding both the rotation angle and flip state, was recorded alongside the match score and location, providing full transparency into the geometric relationship between the two acquisitions. The matched location (*r*^∗^, *c*^∗^) in the scale-normalized EM was mapped back to coordinates in the original high-resolution EM mosaic by inverting the scale factor, yielding the coarse bounding box of the NanoSIMS field of view within the full EM map.

