## Supplementary Figures for "Automated Multimodal Correlative Registration for Organelle-Specific Molecular Imaging"

Lu and Zhao et al.

**Supplementary Figures**

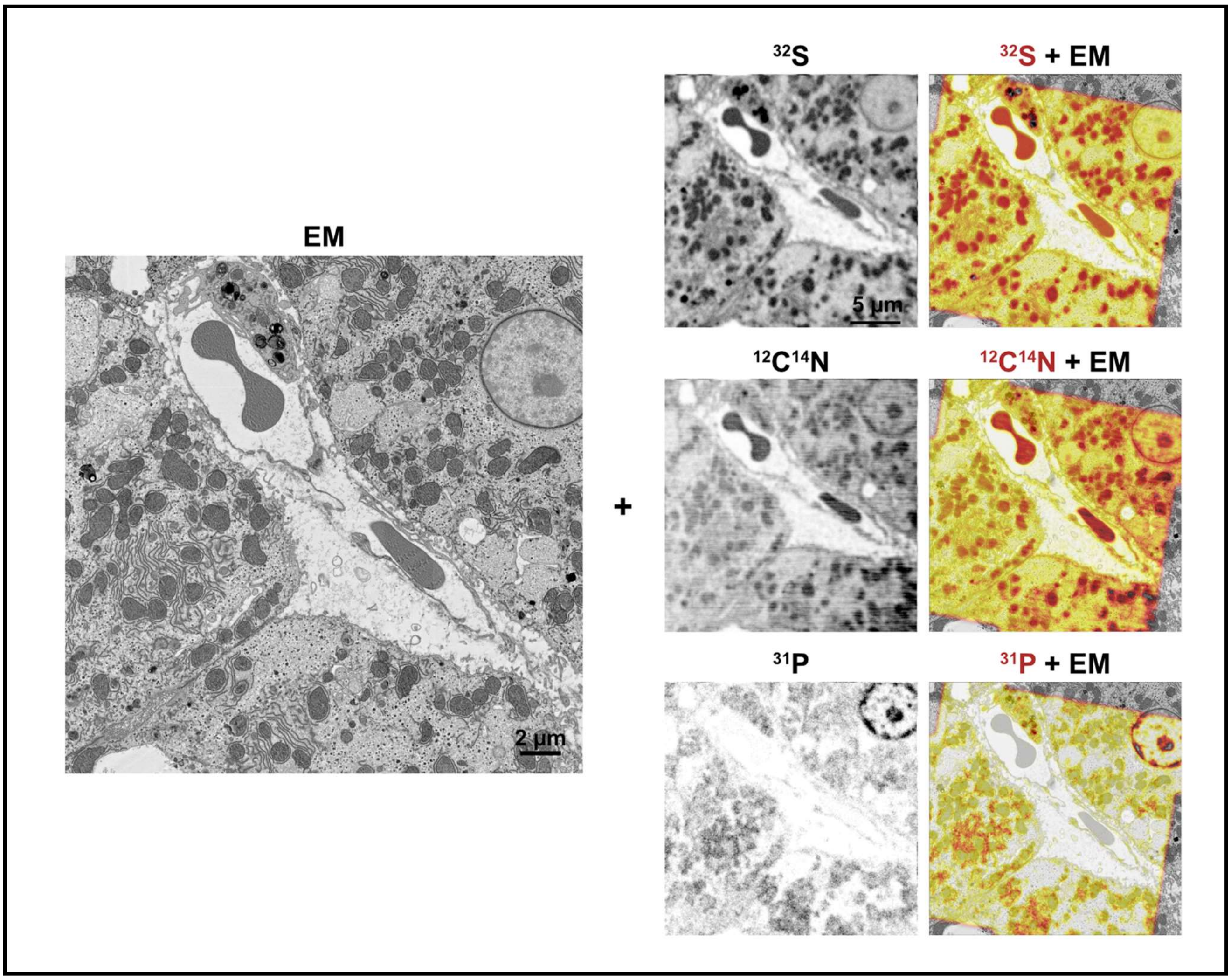


**Fig. S1.** Demonstration of image registration with different secondary ion channels ($^{32}S$, $^{12}C^{14}N$, $^{31}P$). Two panels are shown: a secondary ion image and an overlay of the aligned secondary ion and EM images.

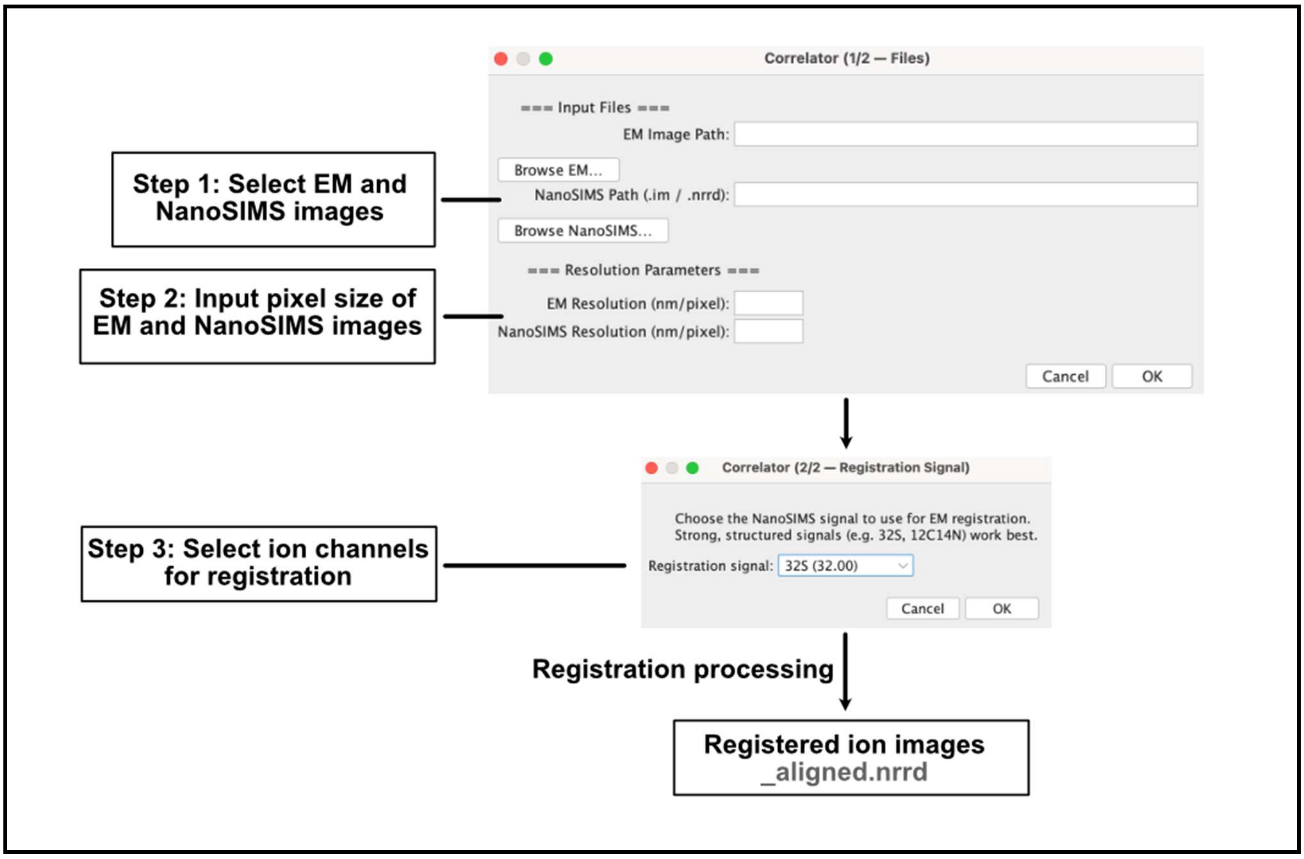


**Fig. S2. Fiji Plugin workflow.** The user browses the EM and NanoSIMS images in the folder and enters the pixel size for both modalities. The channel used for registration is then specified. After processing, the plugin generates an aligned .nrrd file in the same folder as the original NanoSIMS images.

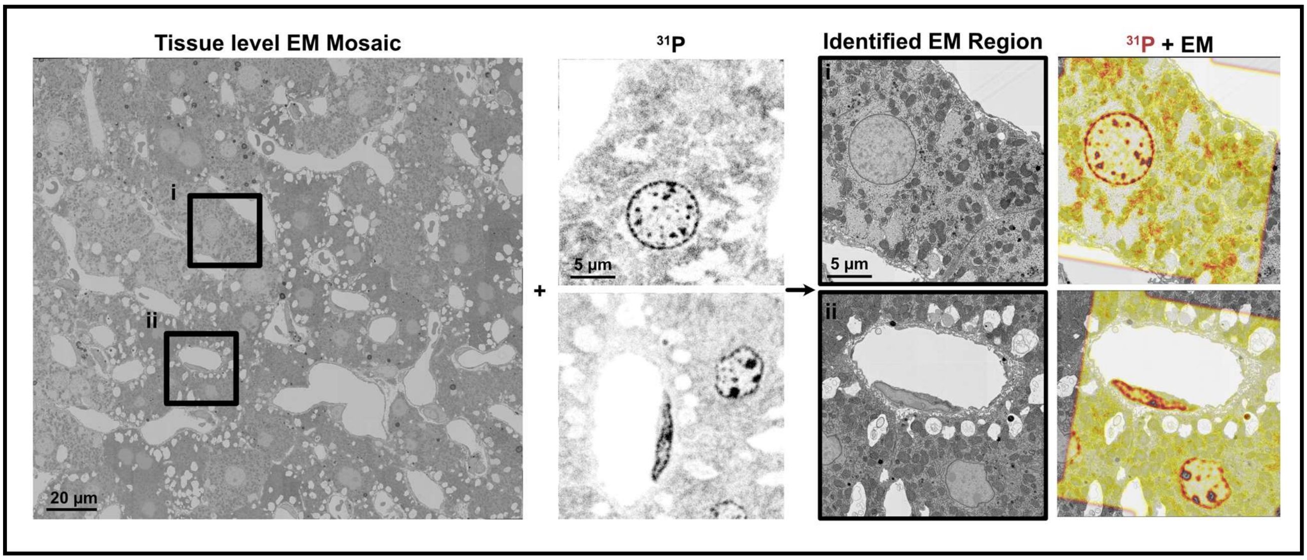


**Fig. S3. Large-field EM alignment via template matching using a $^{31}P$ NanoSIMS image.** Demonstration of alignment of a liver EM mosaic containing two spatially distinct $^{31}P$ image regions (Region i and Region ii) as in Fig. 3b. For each region, $^{31}P$ was used for template matching and registration. The $^{31}P$ image, identified coarse EM patch, and the overlap of $^{31}P$ ion and EM images are shown.

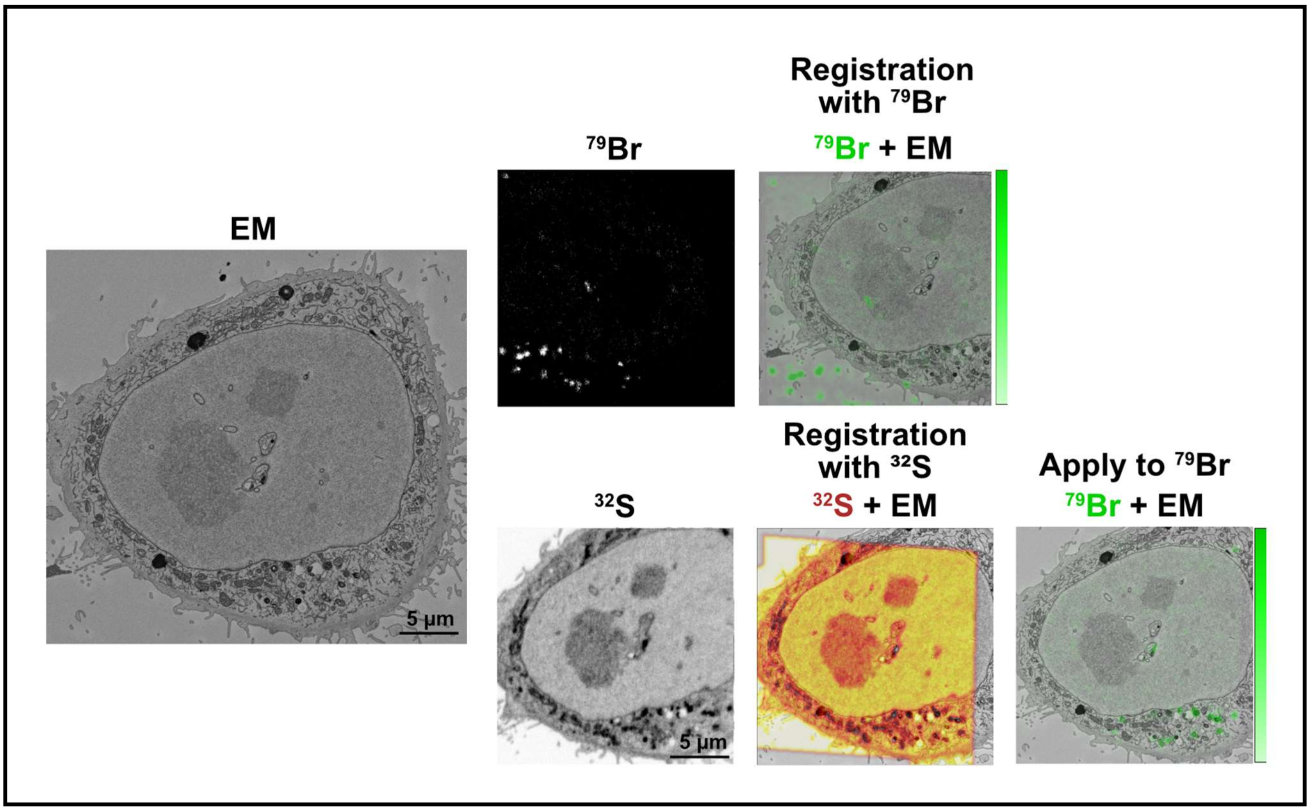


**Fig. S4. Comparison of registration with secondary ion images with and without morphological information.** Demonstration using ion images that are deficient ($^{79}$Br) or rich ($^{32}$S) in morphological information for registration of chemical and ultrastructural images.

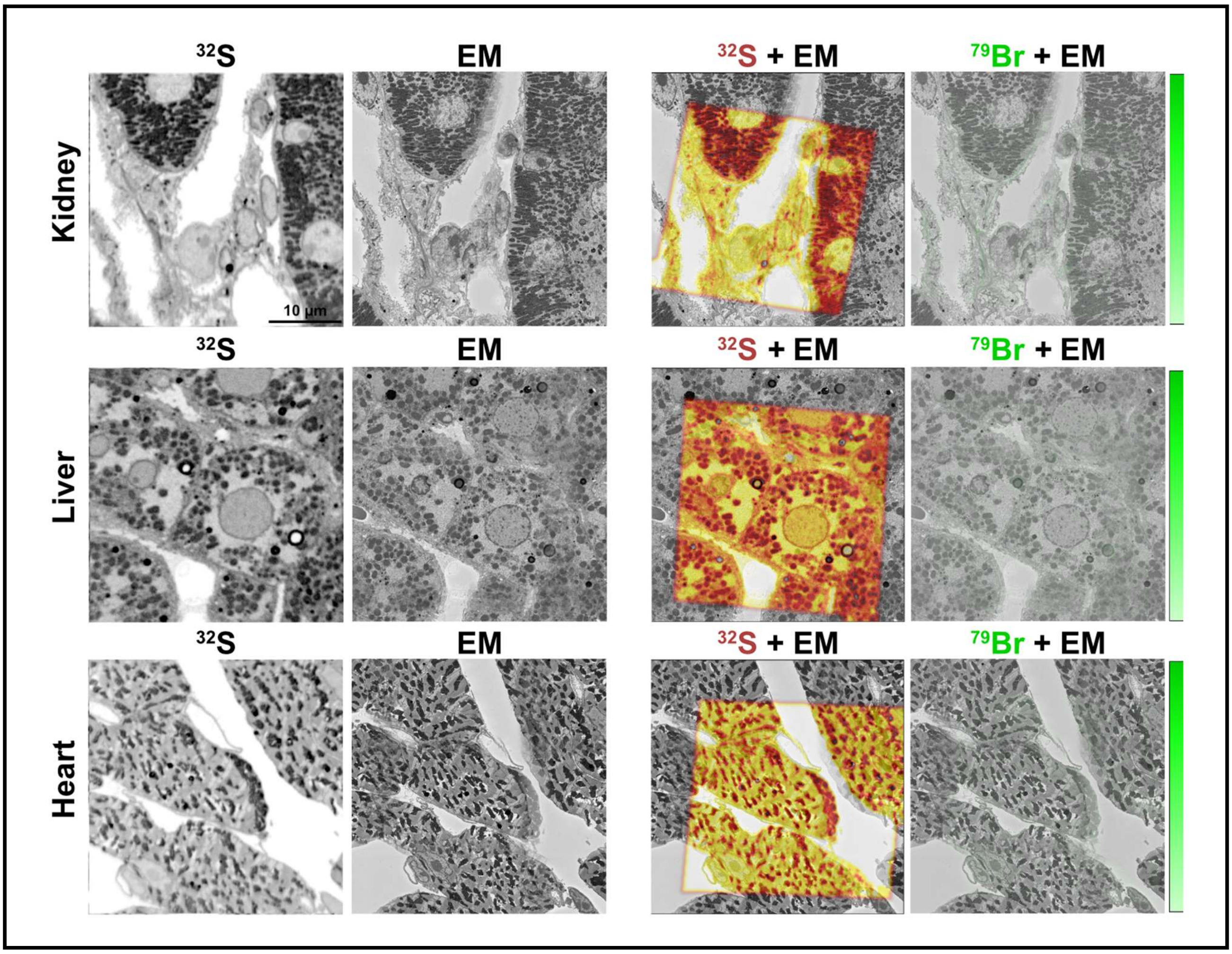


**Fig. S5. Organelle-level mapping of $^{79}$Br signals in kidney, liver, and heart from a control mouse that was not given bromine-labeled ASOs.** Four panels are shown: $^{32}$S image; EM image; overlay of aligned $^{32}$S and EM image; overlay of aligned $^{79}$Br and EM image.

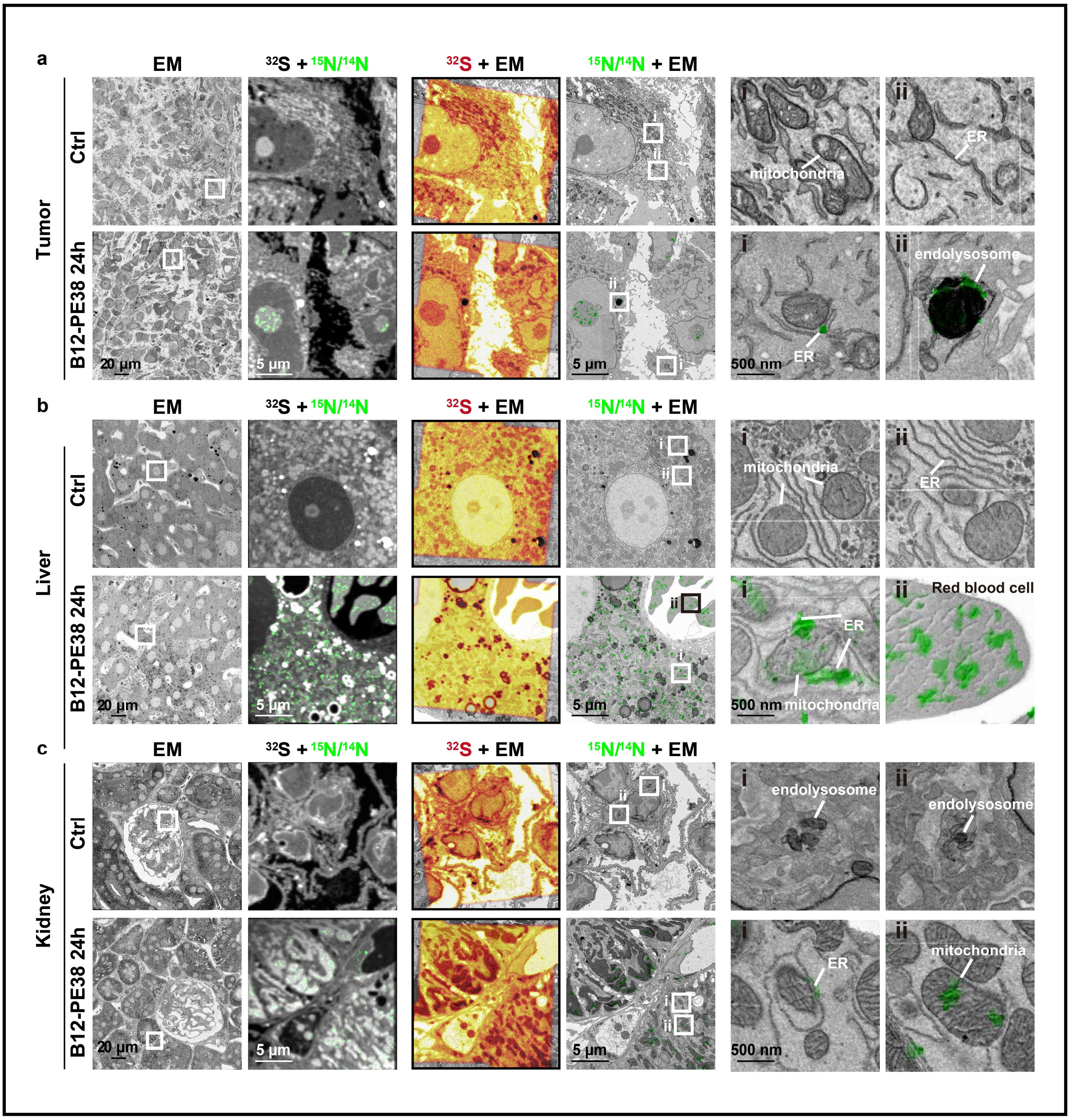


**Fig. S6. Additional examples of organelle-scale mapping of B12-PE38 distribution *in vivo*.** Subcellular mapping of [$^{15}N$] B12-PE38 in tumor (**a**), liver (**b**), and kidney (**c**) tissue 24 h after systemic administration. From left to right: EM image; $^{15}N/^{14}N$ and $^{32}S$ overlay image; $^{32}S$ and EM registered overlay; $^{15}N/^{14}N$ and EM overlay; two representative Zoom-in regions from $^{15}N/^{14}N$ and EM overlays showing organelle-level distributions of B12-PE38 molecules.
